# High-throughput epitope screening of the Human Cytomegalovirus immediate early protein 2 identifies promising antigenic T cell targets

**DOI:** 10.64898/2026.01.24.701419

**Authors:** Eleni Panagioti, Marij J.P. Welters, Ramon Arens, Sjoerd H. van der Burg

**Affiliations:** Department of Surgery, Beth Israel Deaconess Medical Center, Boston, MA, USA; Harvard Medical School, Boston, MA, USA; Department of Medical Oncology, Leiden University Medical Center, Leiden, The Netherlands; Department of Immunology, Leiden University Medical Center, Leiden, The Netherlands

**Keywords:** HCMV, IE2 protein, T cell epitope, Peptide vaccine

## Abstract

Human Cytomegalovirus (HCMV) is an omnipresent pathogen that is associated with increased morbidity and mortality of immunocompromised individuals. Studies of T-cell immunity to HCMV primarily reflect anti-CMV pp65 or immediate early antigen 1 (IE-1) activity. Recent evidence highlights the importance of the major immediate-early 2 (IE2) protein, which is expressed early after HCMV infection and reactivation, for regulating the lytic HCMV replication cycle. In this study, we designed a comprehensive screening approach to assess T cell responses against the IE2 HCMV protein in the peripheral blood of 15 HCMV-seropositive and 6 HCMV-seronegative healthy adults using IE2 synthetic long peptide (SLP) pools and cytokine flow cytometry. The T cell response against the IE2 protein was dominated by CD4^+^ T cells whereas IE2-specific CD8^+^ T-cell reactivity was measured in only 3 donors. Most of the donors recognized chiefly the IE2_351-434_ residues, revealing a remarkably immunogenic area of the protein. Numerous novel HLA class I- and II-restricted IE2 T-cell epitopes were identified. Functional characterization of the IE2 CD4^+^ and CD8^+^ T cell responses uncovered 5 highly antigenic SLPs, which induced polyfunctional Th1 cytokine (IFN-γ^+^/ TNF^+^/ IL-2^+^) response and could serve as candidate vaccine antigens. Evaluation of these 5 highly antigenic IE2 SLPs in T cell-inducing vaccines aiming to inhibit HCMV infection by targeting the expression of immediate-early genes is warranted.

## Introduction

The β-herpesviruses Human Cytomegalovirus (HCMV) is a widespread pathogen that infects a large proportion of the population worldwide^1^. After primary infection, the virus establishes lifelong latency within its host. As a result of productive virus replication immediate early (IE), early and late viral proteins are synthesized. Although, HCMV infection is usually benign, with no clinical disease manifestations in immunocompetent individuals, it can lead to severe complications in susceptible individuals including newborns, allograft recipients, HIV patients and elderly^2–5^.

Despite extensive research, a licensed vaccine intervention to treat or prevent CMV infection remains elusive^6,7^. Studies in patients concur on the importance of T cells for establishment of immunity and long-lasting memory against HCMV^8–15^. Recently, T-cell based vaccines designed to induce CD4^+^ and/or CD8^+^ T-cell responses have received significant research attention. Evidence from various models of mouse CMV infection (MCMV) suggests that vaccine-induced T-cell responses may also successfully control CMV replication^16–20^.

Many of the CMV proteins are recognized by CD4^+^ and CD8^+^ T cells in healthy HCMV-seropositive subjects^21,22^. While T cells most frequently target the antigens pp65 and immediate-early 1 (IE1), the dominance and magnitude of the T-cell response to IE1 is associated with protection from HCMV-induced disease ^23,24^. Interestingly, IE1 but also the major immediate-early 2 (IE2) protein are abundantly expressed and play a key role in initiating lytic cycle virus gene expression and replication^25^. Deletion of the IE1 gene reduces viral replication whereas the IE2 gene is indispensable for virus replication^26–29^. This makes the immediate-early antigens promising HCMV immune targets for vaccine strategies that aim to suppress or clear viral infection at an early phase^30^. While several CMV vaccines against IE1 have been evaluated^31–35^, less is known about the immune response to IE2 protein. Characterization of IE2-specific T-cell immunity in protected healthy individuals will reveal its antigenicity and may foster the development of vaccines targeting indispensable regulatory proteins of HCMV replication.

In this study, we sought to identify novel, HLA class I- and II-restricted T-cell epitopes of the HCMV IE2 protein and enlarge the choice of antigens for potential HCMV T cell-based SLP vaccines. To this end, we mined the immediate early database (IEDB) to map previously identified IE2 epitopes and examined IE2-specific T cell reactivity, directly *ex vivo*, in the peripheral blood of 15 HCMV-seropositive and 6 HCMV-seronegative healthy donors, using overlapping synthetic long peptide (SLP) pools covering the entire IE2 amino acid sequence. Interestingly, IE2-specific T-cell reactivity could be detected directly *ex vivo* in more than half of all subjects screened. The response was dominated by CD4^+^ T cells as IE2-specific CD8^+^ T-cell reactivity was found in only 3 donors. Several previously described but also new HLA class I- and II-restricted IE2 T-cell epitopes were identified. Among all antigenic SLPs there were five, which were frequently recognized among healthy donors, containing epitopes recognized by CD4^+^ T cells and/or CD8^+^ T cells, and considered to be promising candidates for the development of HCMV peptide-based vaccines.

## Methods

### Donors and sample

PBMCs were isolated from buffy coats of 15 HCMV-seropositive and 6 HCMV-seronegative donors following informed consent (Sanquin, The Netherlands) using Ficoll density gradient centrifugation (Ficoll-Amidotrizoaat, Pharmacy LUMC). Cells were cryopreserved in 80% fetal calf serum (FCS; PAA Laboratories) and 20% DMSO (Sigma) and stored in liquid nitrogen at -180 °C. All handling and storage procedures were performed in accordance with the standard operating procedures (SOP) of the Department of Medical Oncology, Leiden University Medical Center.

### Peptides and preparation of peptide pools

The amino acid sequence of HCMV IE2 from the laboratory strain AD169 was used to synthesize 41 synthetic long peptides (SLPs), each 28 amino acids in length and overlapping by 15 residues, collectively covering the entire IE2 protein. Peptide synthesis was performed at the GMP-certified peptide facility of the LUMC. Peptide purity (75-90%) was verified by HPLC, and molecular weight was confirmed by mass spectrometry. In addition, a peptide pool spanning the full pp65 protein and the HLA-A*0201-restricted pp65_495-503_ peptide (NLVPMVATV) were included in T cell detection assays. Peptides were dissolved in DMSO at a stock concentration of 50 mg/mL, further diluted to 2 mg/mL, and stored at -20 °C. Eight IE2 peptide pools, each comprising 5-6 SLPs, were subsequently prepared and stored under the same conditions. All individual IE2 peptides and peptide pools used in this study are listed in Supplementary Table 1.

### Ex vivo detection of antigen-specific T cells

Direct *ex vivo* detection of HCMV-specific T cells was performed as previously described^36^. Briefly, autologous plastic-adherent monocytes were stimulated with granulocyte–macrophage colony-stimulating factor (GM-CSF; 800 U/mL; Invitrogen) for 2-4 days at 37 °C in 5% CO₂, after which they were incubated for 5 h with individual IE2 peptide pools, a pool of all IE2 peptides, a pool of all pp65 peptides, or the pp65 short peptide (SP) at a final concentration of 50 µg/mL, in the presence of poly (I:C) (25 µg/mL; Invivogen). Medium alone served as a negative control, and Staphylococcal enterotoxin B (SEB; 1 µg/mL; Sigma) was used as a positive control. Autologous PBMCs (2-4 × 10⁶ cells/well) and Roferon (interferon alpha 2a; 3 × 10⁶ U/0.5 mL; Roche) were added to the monocyte cultures. One hour later, Brefeldin A (Sigma) was added, and cells were incubated for 16-20 h at 37 °C to allow accumulation of intracellular cytokines. Following incubation, T cells were stained with a live/dead marker (Yellow ARD; Life Technologies) and antibodies against CD3 (UCHT1), CD4 (SK3), CD8 (SK1), CD14 (M5E2), IFN-γ (B27), IL-2 (5344.111), CD137 (4B4-1), CD154 (TRAP1) from BD Biosciences, and TNF (MAb11) and CD45RA (HI100) from BioLegend, following standard operating procedures of the Department of Medical Oncology. Samples were acquired on an LSRFortessa cytometer (BD Biosciences) and analyzed using FlowJo v10 software (Tree Star). Flow cytometry gating strategies are shown in Supplementary Figure 1. An immune response was considered positive when donor T cells co-produced IFN-γ and TNF at levels ≥2-fold above background (medium control).

### In vitro expansion of low frequency antigen-specific T cells

To validate *ex vivo* detected T cell responses to IE2 peptide pools and map reactivity to individual SLPs, PBMCs were subjected to a 10-day antigen-specific T cell stimulation culture^37^. Briefly, donor PBMCs were thawed, resuspended in IMDM supplemented with 10% HAB, and seeded in triplicate (3 mL/well) in 6-well plates (Costar). Peptide pools were added at a final concentration of 2.5 µg/mL, and cells were cultured overnight at 37 °C in 5% CO₂. The following day, IL-15 (5 ng/mL; Peprotech) and T cell growth factor (TCGF; Zepto Metrix) at 10% final concentration were added, and cultures were maintained for 10 days (referred to as bulk cultures). In parallel, on day 7, autologous PBMCs were thawed, seeded in 24-well plates, and monocytes were cultured for 2 days at 37 °C. On day 9, adherent monocytes were loaded with the respective peptide pools and individual SLPs at 5 µg/mL. On day 10, bulk-cultured T cells (7 × 10⁵-1 × 10⁶ cells/mL) were added to peptide-loaded monocytes. After 1 h, Brefeldin A was added, and cells were incubated for 16-20 h at 37 °C. Positive and negative controls identical to those used in the *ex vivo* assay were included. Intracellular cytokine staining (ICS) was performed, and samples were acquired and analyzed as described above. Responses were considered positive when donor T cells produced IFN-γ at levels ≥2-fold above background (medium control).

### Statistical analysis

Statistical comparisons were performed using unpaired Student’s t-tests or one-way ANOVA in GraphPad Prism v6 (GraphPad Software, USA). Significance was defined as *p* < 0.05, \**p* < 0.01, and \*\**p* < 0.001.

## Results

### Ex vivo detection of low-frequency CD4⁺ and CD8⁺ T cell responses to HCMV IE2 peptide pools

To identify novel HCMV IE2-derived T cell epitopes restricted by HLA class I and class II molecules, we employed a traditional, high-throughput and well-established screening approach using overlapping synthetic long peptides (SLPs)^36^. Notably, the use of overlapping SLPs facilitates rapid identification of the precise peptide sequences recognized upon detection of a positive response. Synoptic illustration of the screening strategy is presented in **Figure 1**. A schematic overview of the screening strategy is shown in **Figure 1**. A cohort of 21 healthy volunteers (15 CMV-seropositive and 6 CMV-seronegative), aged 34-65 years, was recruited irrespective of gender, ethnicity or educational background (**Table 1**).

**Figure 1.**
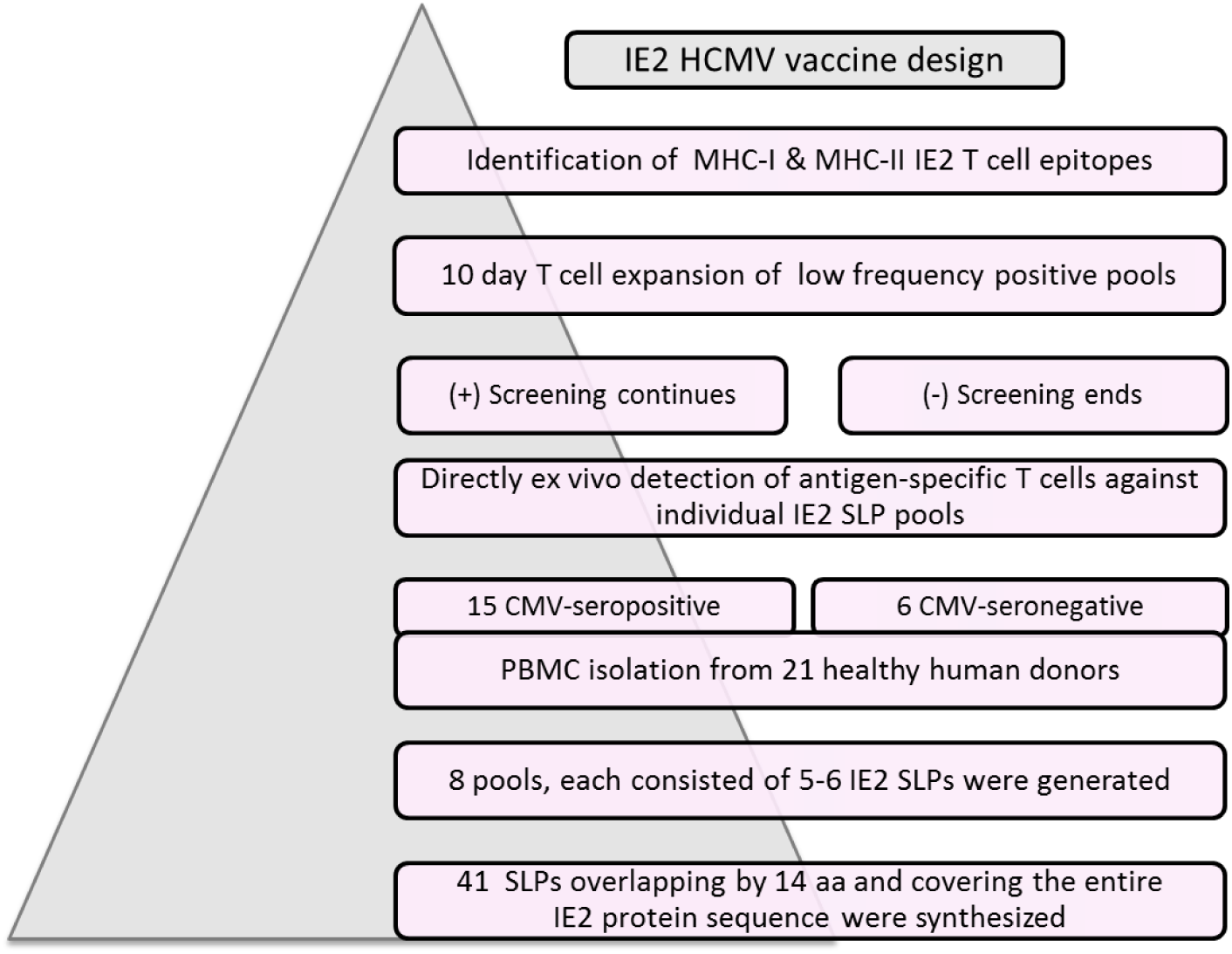
Overview of the IE2 T cell epitope screening strategy. Forty-one synthetic long peptides (SLPs, 28 aa each, overlapping by 14 aa) covering the entire HCMV IE2 protein were synthesized and organized into eight pools, each containing 5-6 SLPs. PBMCs from 21 healthy donors (15 CMV-seropositive, 6 CMV-seronegative) were screened by *ex vivo* intracellular cytokine staining (ICS) for direct recognition of IE2 peptide pools. Low-frequency peptide-reactive T cells were expanded *in vitro* for 10 days, and subsequently analyzed by ICS and flow cytometry to assess CD4⁺ and CD8⁺ T cell responses. Positive responses to individual SLPs were confirmed, enabling identification of IE2-specific HLA class I and II epitopes. Five frequently recognized IE2 peptides were identified, representing potential candidates for inclusion in an HCMV vaccine.

**Table 1.** Donor demographics and summary of *ex vivo* pp65 and IE2 T cell responses. PBMCs from 21 healthy donors were screened for CD4⁺ and CD8⁺ T cell recognition of HCMV pp65 and IE2 peptide pools. Donor characteristics including ID number, age, gender (M, male; F, female), and CMV serostatus are indicated. Available HLA-A/B/C and HLA-DR/DQ/DP types are shown. Ex vivo CD4⁺ and CD8⁺ T cell responses against IE2 and pp65 pools are summarized as positive (+) or negative (-) for each donor.

|  |  |  |  | Ex vivo T cell response |  | Class I |  |  | Class II |  |
| --- | --- | --- | --- | --- | --- | --- | --- | --- | --- | --- |
| Donor # | Age | Gender | CMV status | pp65<br>CD4/CD8 | IE2<br>CD4/CD8 | HLA-A | HLA-B | HLA-C | HLA-DR | HLA-DQ |
| 1 | 65 | M | + | +/+ | +/- | 2, 11 | 55, 35 | 3, 4 | 1, 15 | 5, 6 |
| 2 | 56 | M | + | +/+ | -/- | 2, 3 | 44, 6 | 3, - |  |  |
| 3 | 62 | M | + | -/+ | +/- | 2, 3 | 27, 35 | 2, 4 | 1, 4 |  |
| 4 | 48 | M | + | -/- | -/- | 2, 3 | 7, 35 | 4, 7 | 1, 15 | 1 |
| 5 | 53 | F | + | -/- | +/- | 2, 3 | 7, 35 | 4, 7 |  |  |
| 6 | 46 | F | + | +/+ | +/- |  |  |  |  |  |
| 7 | 64 | F | + | +/+ | +/- | 1, 2 | 44, 56 |  |  |  |
| 8 | 51 | F | + | +/- | +/- |  |  |  |  |  |
| 9 | 56 | F | + | -/- | -/- | 2, 29 | 7, 44 | 7 | 4, 7 | 2, 8 |
| 10 | 51 | M | + | +/+ | +/+ |  |  |  |  |  |
| 11 | 42 | M | + | +/+ | +/- |  |  |  |  |  |
| 12 | 44 | F | + | +/+ | +/- | 2, 24 | 62, 38 | 3 | 4, 14 | 5, 3 |
| 13 | 45 | M | + | -/- | -/- |  |  |  |  |  |
| 14 | 61 | M | + | -/+ | -/- |  |  |  |  |  |
| 15 | 55 | F | + | -/- | -/- | 2, 3 | 44, 62 | 3 | 4, 7 | 2, 3 |
| 16 | 54 | M | - | -/- | +/- | 31, 68 | 51, 39 |  | 11, 14 | 5, 7 |
| 17 | 34 | M | - | -/- | -/- |  |  |  |  |  |
| 18 | 54 | M | - | -/- | -/- | 1, 3 | 39, 35 | 4 |  |  |
| 19 | 53 | F | - | -/- | -/- |  |  |  |  |  |
| 20 | 46 | M | - | -/- | +/- | 23, 68 | 51, 44 | 4 |  |  |
| 21 | 58 | M | - | -/- | -/- | 2, 24 | 14, 14 | 6 |  |  |

To assess virus-primed T-cell reactivity, the directly *ex vivo* detectable IE2-specific cellular responses to each peptide pool were quantified based on the frequency of antigen-specific IFN-γ and TNF-producing CD4^+^ and CD8^+^ T cells. IE2-specific CD4^+^ T cell responses were detected in 11 of 15 CMV-seropositive donors, whereas IE2-specific CD8^+^ T cell responses were observed in 3 of these individuals (**Table 1**, **Figure 2A and B**). Notably, IE2-specific CD4^+^ T cell reactivity was also detected in 3 CMV-seronegative donors (donors 16, 17 and 20). Peptide pool 6 (IE2_351-434_) was the most frequently recognized antigenic region (9 of 2 responders), followed by pool 1 (IE2_1-84_) and pool 7 (IE2_421-504_). In contrast, peptide pools 3, 4 and 5 (IE2_141-364_) elicited minimal responses (**Figure 2A and B**). Overall, the magnitude of *ex vivo* detected responses was low, limiting detailed phenotypic characterization. Representative examples of IFN-γ⁺ TNF⁺ CD4⁺ T cell responses from CMV-seropositive donors (donors 1 and 6) and a CMV-seronegative donor (donor 16) following stimulation with IE2 peptide pools are shown in **Figure 3**.

**Figure 2.**
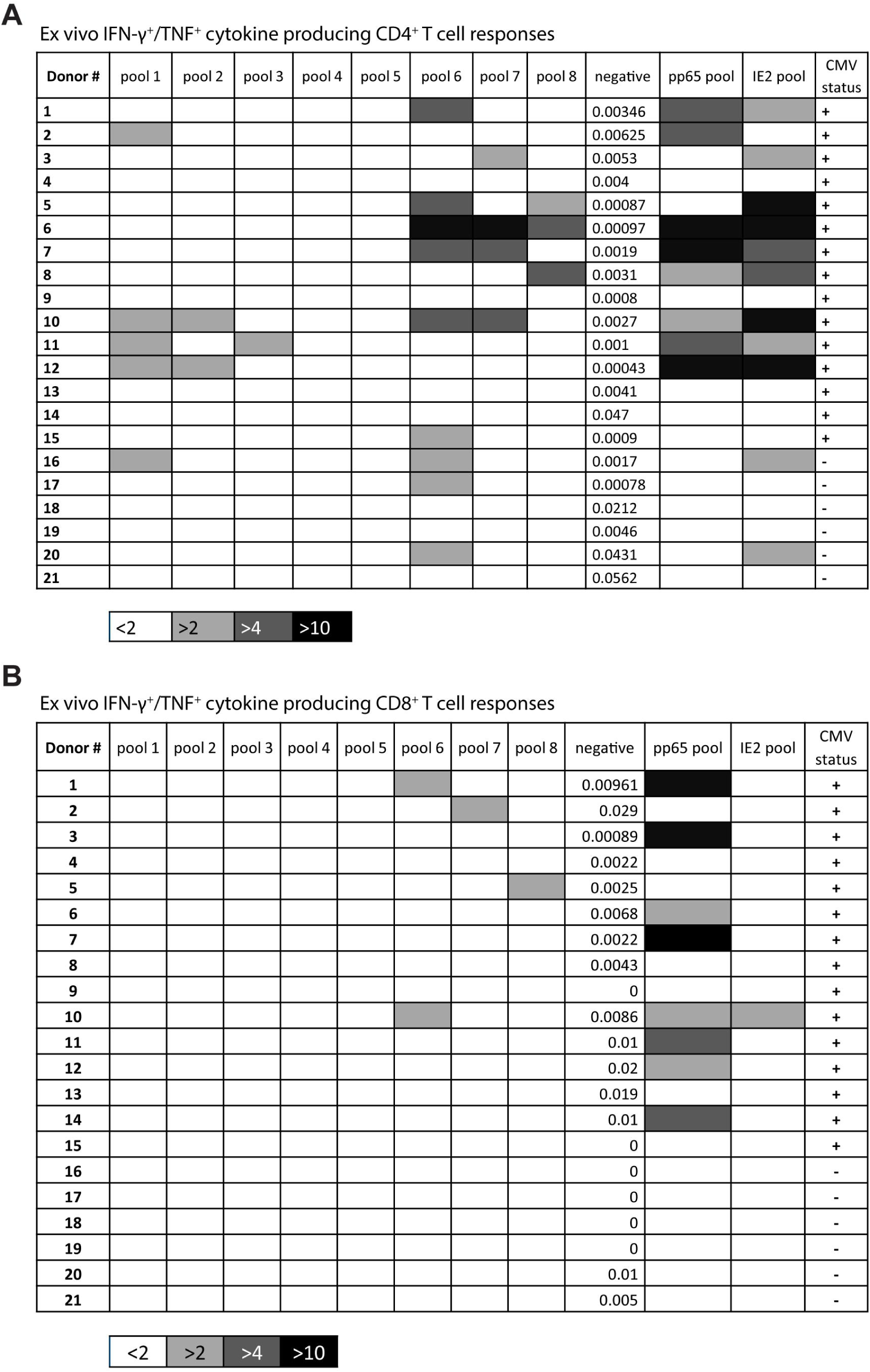
Ex vivo CD4⁺ and CD8⁺ T cell recognition of IE2 and pp65 peptide pools. Ex vivo IFN-γ⁺/TNF⁺ CD4⁺ **(A)** and CD8⁺ **(B)** T cell responses were measured in each donor following stimulation with IE2 and pp65 peptide pools. Donor number and CMV serostatus are indicated. Responses in unstimulated samples represent background levels. Cellular responses ≤2-fold above background (white boxes) were considered negative, whereas responses >2-fold above background (light grey boxes) were considered positive and selected for further evaluation. Positive CD4⁺ T cell responses were detected in 14 donors, and positive CD8⁺ T cell responses were observed in 3 donors.

**Figure 3.**
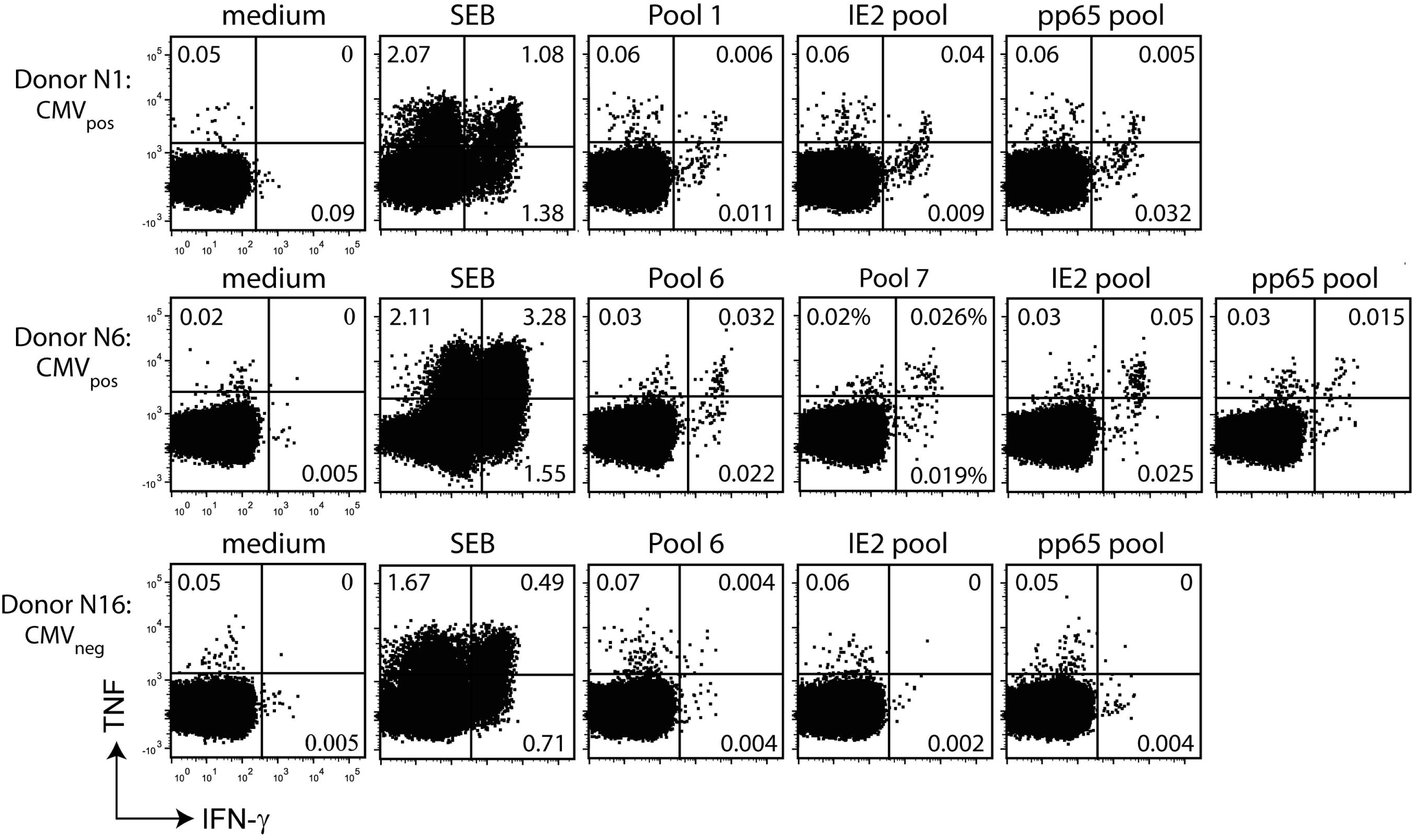
CD4⁺ T cell recognition of IE2 peptide pools in HCMV-seropositive and -seronegative donors. PBMCs from donors were stimulated with eight IE2 peptide pools (5-6 SLPs per pool), and T cell reactivity was assessed by ICS and flow cytometry. Representative *ex vivo* IFN-γ⁺/TNF⁺ CD4⁺ T cell responses are shown for CMV-seropositive donors (donors 1 and 6) and a CMV-seronegative donor (donor 16). Background responses were measured in unstimulated PBMCs (medium), and SEB was used as a positive control. In the fluorescence plots, the x-axis represents IFN-γ expression and the y-axis TNF production within CD4⁺ T cells.

In parallel, T cell responses to a pp65 peptide pool were assessed to enable comparison of the magnitude and immunological relevance of IE2-specific T cell responses during HCMV infection. Although response levels varied among donors, most CMV-seropositive individuals exhibited CD4⁺ T cell reactivity to both pp65 and IE2 antigens (**Table 1**, **Figure 2 and Figure 4A**). However, the magnitude of pp65-specific CD4⁺ T cell responses did not directly correlate with responses to the corresponding IE2 peptide pools (**Figure 4B**). Moreover, whereas IE2-specific CD8⁺ T cell responses were infrequently detected, the pp65 SLP pool elicited CD8⁺ T cell reactivity in 8 of 15 CMV-seropositive (**Figure 4C**, **Table 1 and Figure 2B**).

**Figure 4.**
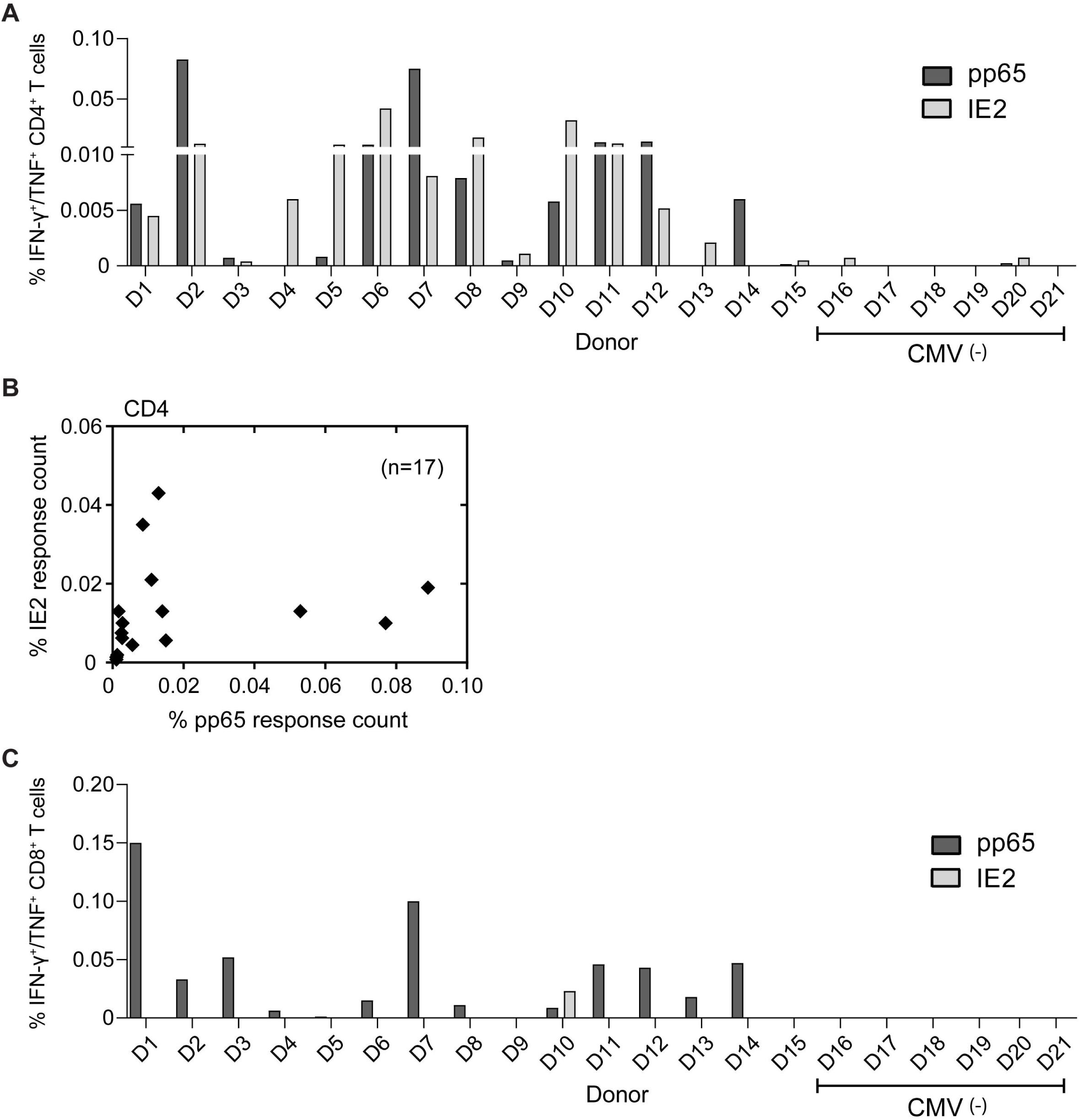
Characterization of total IE2- and pp65-specific CD4⁺ and CD8⁺ T cell responses in HCMV-seropositive and -seronegative donors. PBMCs from 21 donors (15 CMV-seropositive, 6 CMV-seronegative) were screened by *ex vivo* ICS to assess direct CD4⁺ and CD8⁺ T cell reactivity against IE2 and pp65 peptide pools. **(A)** Percentages of total IFN-γ⁺/TNF⁺ CD4⁺ T cell responses to pp65 and IE2 pools across the donor cohort. **(B)** No correlation was observed between the magnitude of pp65 and IE2 CD4⁺ T cell responses among responder donors (n = 12; *p* > 0.055). **(C)** Frequency of IFN-γ⁺/TNF⁺ CD8⁺ T cell responses against pp65 and IE2 pools. IE2-specific CD8⁺ T cell reactivity was detected only in donor 10, whereas pp65-specific CD8⁺ responses were observed in 8 of 15 CMV-seropositive donors.

Collectively, *ex vivo* immune reactivity analyses indicate that the HCMV IE2 protein has a strong propensity to elicit CD4⁺ T cell responses, comparable to those induced by pp65. In particular, the region encompassed by IE2 peptide pool 6 (IE2_351-434_) was recognized by the majority of donors, identifying a highly immunogenic domain within the protein. By contrast, and unlike pp65, IE2 did not efficiently induce *ex vivo* detectable CD8⁺ T cell responses.

### Identification of HLA class I- and class II-restricted IE2 T cell epitopes

To validate the *ex vivo* detected IE2-specific T cell responses and to identify the cognate peptides within each reactive IE2 peptide pool, PBMCs from all responding donors were stimulated with individual peptides. Antigen-specific T cells were expanded *in vitro* for 10 days, after which cells were restimulated with each synthetic long peptide (SLP) contained within the corresponding pool and analyzed by polychromatic intracellular cytokine staining (ICS). Peptide-specific CD4⁺ and CD8⁺ T cell responses were detected in 11 and 3 donors, respectively, confirming the *ex vivo* findings. Representative examples of robust peptide-specific T cell reactivity observed in expanded PBMCs from both CMV-seropositive and CMV-seronegative donors are shown in **Figure 5**. While some donors responded to a single SLP, most recognized two or more peptides, with donor 6 exhibiting CD4⁺ T cell reactivity to 13 distinct SLPs **(Supplementary Table 2)**. Overall, *in vitro* expansion revealed a substantially broader repertoire of IE2-specific T cell epitopes than was apparent from *ex vivo* ICS using peptide pools. A summary of all CD4⁺ and CD8⁺ T cell responses detected against individual SLPs is provided in **Tables 2 and 3**, respectively.

**Figure 5.**
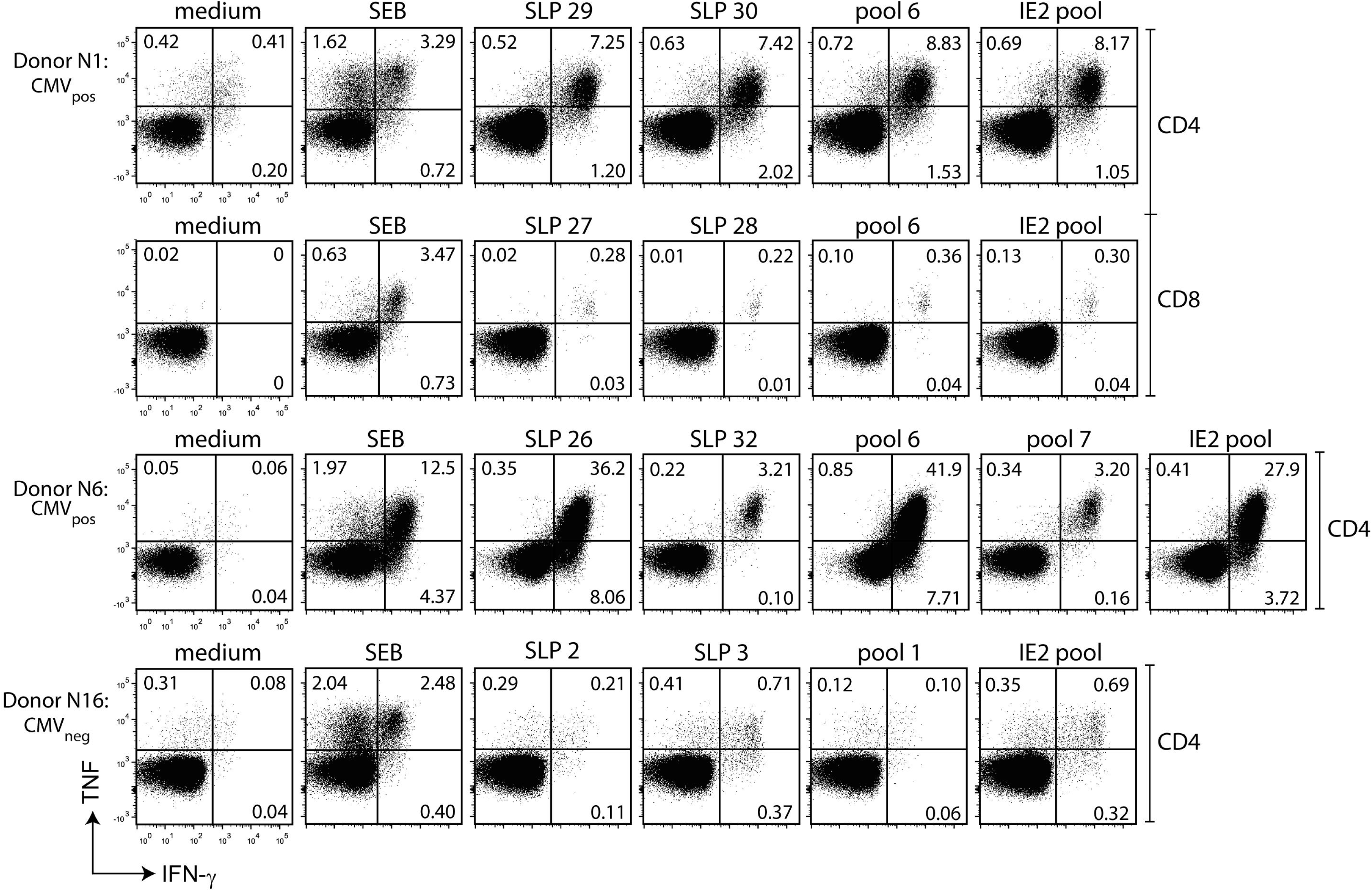
CD4⁺ and CD8⁺ T cell epitope identification in HCMV-seropositive and -seronegative donors. PBMCs that responded to IE2 peptide pools *ex vivo* were expanded *in vitro* for 10 days and subsequently restimulated with individual SLPs from the corresponding pools. Representative ICS plots are shown for two CMV-seropositive donors (donors 1 and 6) and one CMV-seronegative donor (donor 16). In donor 1, CD4⁺ and CD8⁺ T cell reactivity to SLPs 29 and 30 indicates the presence of HLA class II and class I epitopes, respectively. Responses to individual SLPs, peptide pools, and the full IE2 pool are shown. Unstimulated PBMCs (medium) and SEB served as negative and positive controls, respectively. In the fluorescence plots, the x-axis represents IFN-γ expression and the y-axis TNF production within CD4⁺ or CD8⁺ T cells.

**Table 2.**
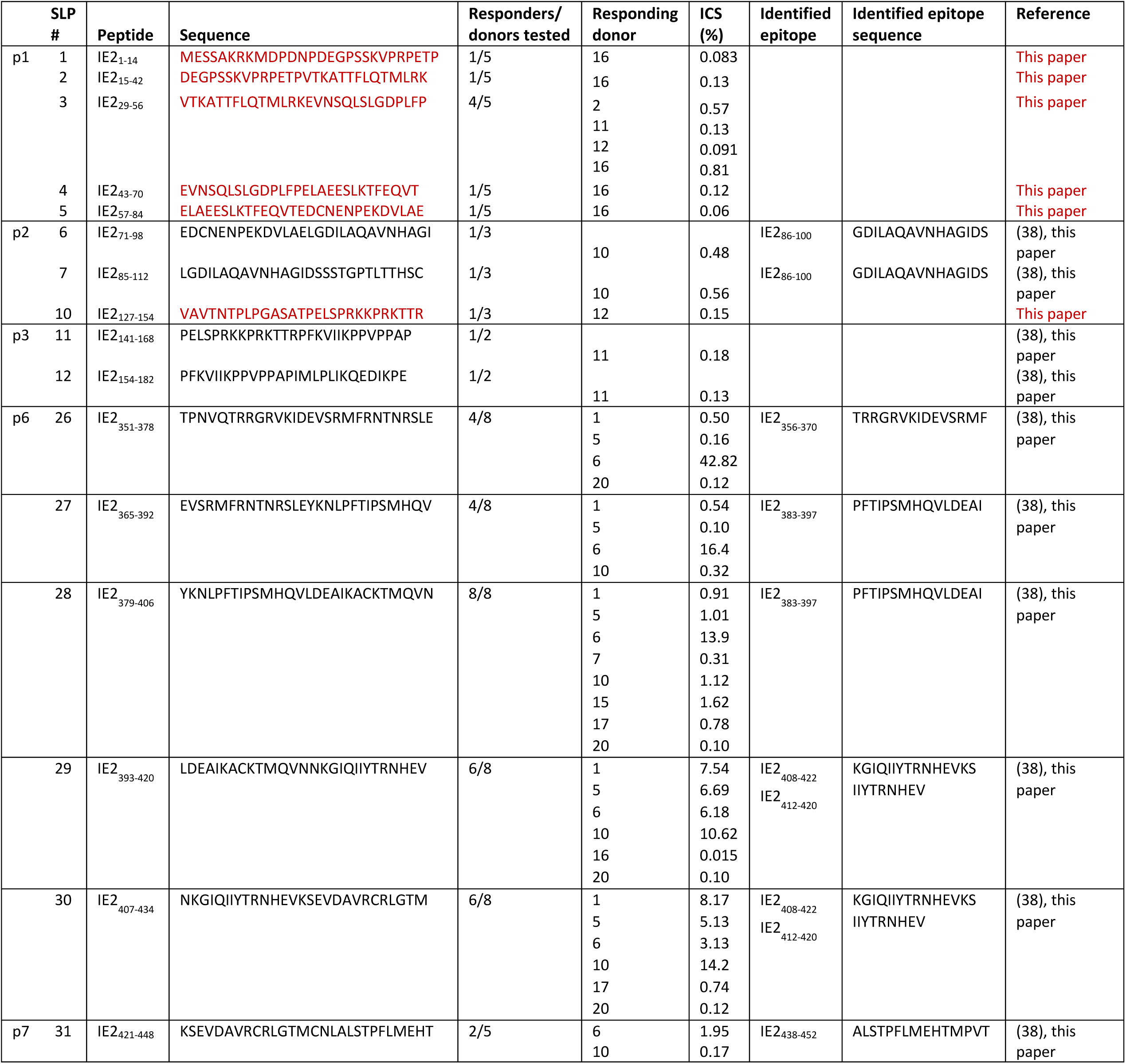
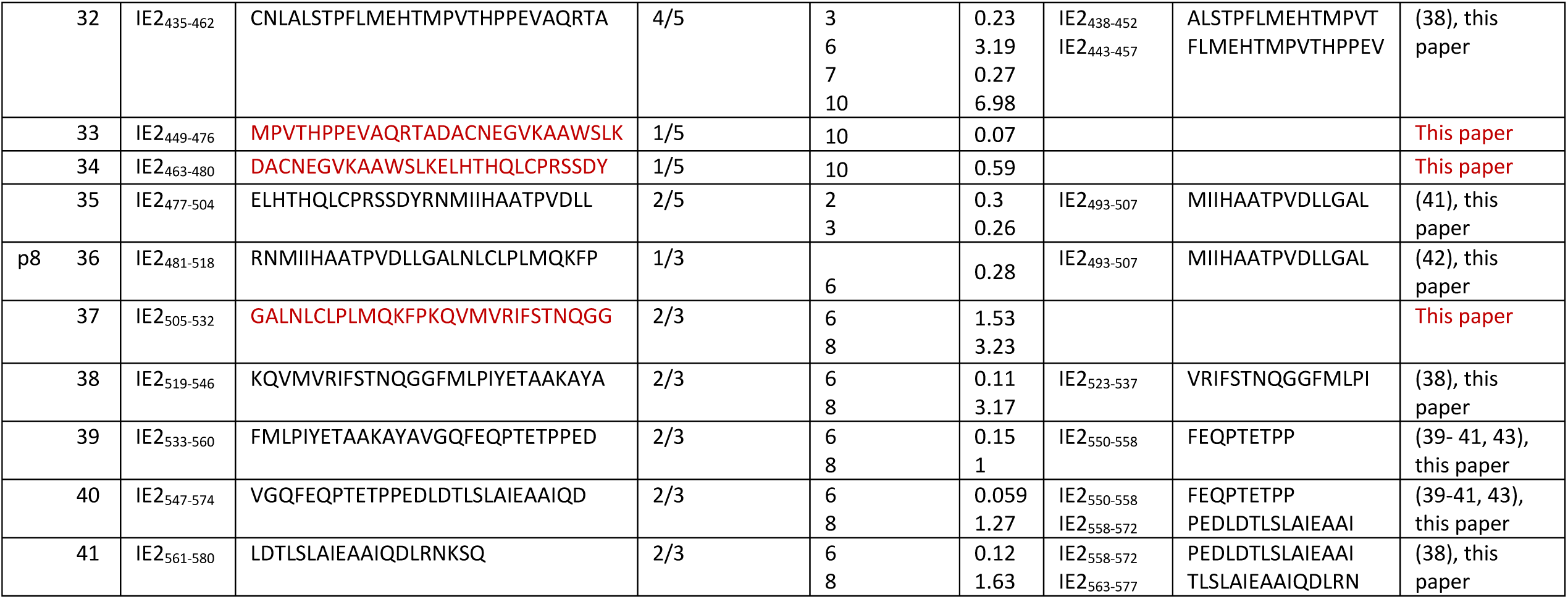
Identified IE2-specific CD4⁺ T cell epitopes. CD4⁺ T cell responses to individual IE2 peptide pools (pool 1 = p1) and the number of SLPs per pool are shown. SLP amino acid positions and sequences are listed. The number of responding donors and the frequency of IFN-γ⁺ CD4⁺ T cells measured by ICS are indicated. Novel epitopes identified in this study are highlighted in red, and previously reported epitopes are shown in black.

**Table 3.** Identified IE2-specific CD8⁺ T cell epitopes. Characteristics of detected CD8⁺ T cell responses are shown. Two IE2 MHC class I epitopes were identified. Submer peptides for all detected epitopes were synthesized, and their recognition was validated by ICS. A previously reported epitope (IE2_42-50_) was confirmed, and a novel epitope (IE2_382-390_) is highlighted in red.

| SLP # | Peptide | Sequence | Responder/<br>donors<br>tested | Responding<br>donors | ICS<br>(%) | Identified<br>epitope | Identified<br>epitope<br>sequence | Reference |
| --- | --- | --- | --- | --- | --- | --- | --- | --- |
| p1 3 | IE2<sub>29-56</sub> | VTKATTFLQTMLRKEVNSQLSLGDPLFP | 1/5 | 2 | 0.38 | IE2<sub>42-50</sub> | KEVNSQLSL | (38, 44-47), this paper |
| p6 27 | IE2<sub>365-392</sub> | EVSRMFRNTNRSLEYKNLPFTIPSMHQV | 2/7 | 1<br>10 | 0.29<br>0.029 | IE2<sub>82-90</sub> | LPFTIPSMH | This paper |
| 28 | IE2<sub>379-406</sub> | YKNLPFTIPSMHQVLDEAIKACKTMQVN | 2/7 | 1<br>10 | 0.23<br>0.02 |  |  |  |
| p8 38 | IE2<sub>519-546</sub> | KQVMVRIFSTNQGGFMLPIYETAAKAYA | 1/3 | 5 | 0.086 |  |  | This paper |
| 39 | IE2<sub>533-560</sub> | FMLPIYETAAKAYAVGQFEQTETPPED | 1/3 | 5 | 0.071 |  |  |  |

To contextualize these findings, the Immune Epitope Database (IEDB) was queried for all previously reported HLA class I- and class II-restricted T cell epitopes derived from the HCMV IE1 (UL123) and IE2 (UL122) proteins up to May 2023. A schematic representation of the epitope locations and their HLA restrictions, including those identified in the present study, is shown in **Supplementary Figure 2**. Notably, most previously reported IE2-specific HLA class I- and class II-restricted T cell epitopes were confirmed^38–47^, and, in addition, six novel CD4⁺ T cell epitopes and one novel CD8⁺ T cell epitope were identified.

### Functional profiling of T cell reactivity against immunodominant IE2 epitopes

The frequency of responding individuals serves as an indicator of peptide antigenicity^48^. Among the 41 individual IE2 synthetic long peptides (SLPs) tested, five contained unique sequences that were recognized by CD4⁺ T cells (SLPs 3, 26, 28, 30 and 32) or CD8⁺ T cells (SLPs 27/28) in the majority of donors (peptide sequences highlighted in bold in **Tables 2 and 3**). Because the functional quality of peptide-specific T cell responses is a critical determinant of their potential protective efficacy against CMV infection, we next assessed the cytokine profiles elicited by IE2-specific T cell epitopes. All selected peptides induced robust, polyfunctional responses, as evidenced by the proportion of T cells co-producing the antiviral cytokines IFN-γ, TNF and IL-2. Collectively, these analyses identified five highly antigenic IE2 SLPs capable of eliciting polyfunctional type 1 T cell responses (**Figure 6A-F**).

**Figure 6.**
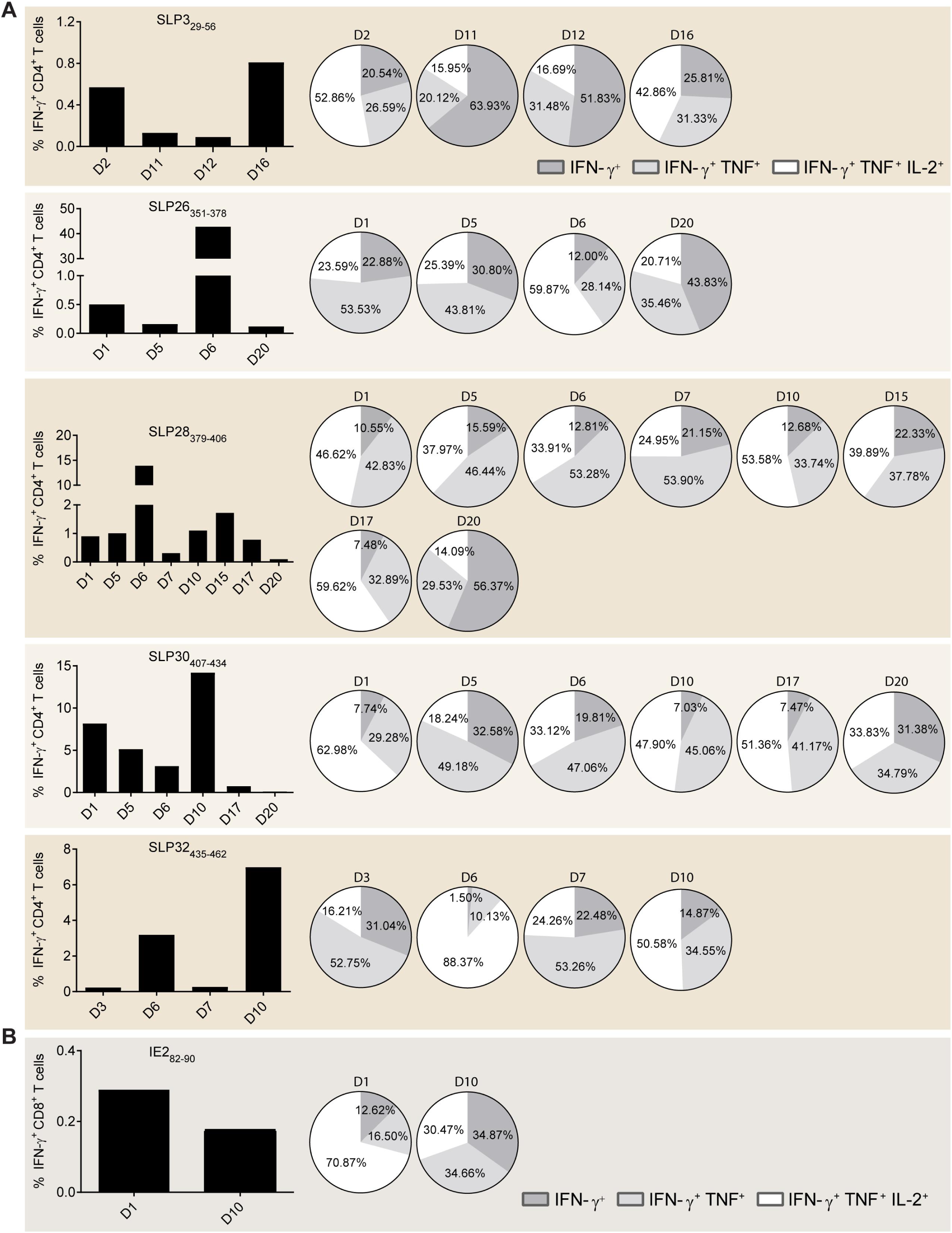
Functional profiling of the most immunogenic HLA class I and II IE2 T cell epitopes. **(A-E)** Percentages of IFN-γ⁺ CD4⁺ T cells and **(F)** CD8⁺ T cells from responder donors following restimulation with the respective peptide epitopes. Cytokine production was measured in *in vitro* expanded T cells using polychromatic ICS. Pie charts indicate the proportions of single (IFN-γ), double (IFN-γ/TNF), and triple (IFN-γ/TNF/IL-2) cytokine-producing cells within each antigen-specific CD4⁺ or CD8⁺ T cell population across all responding donors.

## Discussion

Attacking viral infection at an early stage may prevent widespread dissemination and full-blown disease. Reinforcing T cell responses against highly immunogenic antigens expressed immediately after infection could therefore be critical for the development of protective vaccines. The HCMV IE2 protein, expressed during the immediate-early phase of infection and reactivation, represents a particularly valuable target for such strategies. In this study, we systematically evaluated the immunogenicity of IE2 to identify amino acid stretches frequently recognized by T cells from healthy, HCMV-seropositive donors. This analysis revealed numerous MHC class I- and II-restricted epitopes, including both previously reported^38–47^ and novel sequences, detectable directly *ex vivo*. The majority of immunodominant epitopes mapped to residues IE2_351-434_, highlighting this region as a hotspot for T cell recognition. Furthermore, five SLPs were consistently recognized across donors expressing diverse HLA alleles. These SLPs contained epitopes recognized by CD4⁺ and/or CD8^+^ T cells^38,44–47^ and represent attractive candidates for inclusion in an IE2-targeted T cell–based vaccine.

Unexpectedly, IE2-specific CD4⁺ T cell responses were detected *ex vivo* and after *in vitro* expansion in three of six CMV-seronegative donors. Similar phenomena have been reported in other viral infections; for example, HIV-specific memory T cells have been observed in highly exposed, seronegative individuals^49,50^. This finding suggests that CMV exposure does not invariably result in persistent infection and that heterogeneity in susceptibility may reflect natural protective immunity. Factors such as the size of the initial infectious inoculum could contribute to these differences, influencing both humoral and T cell responses^51,52^. Although our study could not define the activation or memory status of these low-frequency responses, functional characterization in larger cohorts may reveal biomarkers of natural resistance.

Analysis of IE2-specific responses revealed a predominance of *ex vivo* detectable CD4⁺ T cell activity (11 of 15 seropositive donors), whereas CD8⁺ T cell responses were observed in only three donors. This pattern is consistent with previous studies^38^ and cannot be attributed to technical limitations, as pp65-specific CD8⁺ T cell responses were readily detected in parallel. Robust induction of both CD4⁺ and CD8⁺ T cell responses with broad functional capacity is considered essential for vaccine efficacy^53,54^. Indeed, murine studies using SLP vaccines incorporating both MHC class I- and II-restricted epitopes from multiple immunogenic proteins achieved optimal containment of MCMV upon challenge^16,18,20^. These findings support the translational potential of identifying immunodominant epitopes in human HCMV proteins. Interestingly, whereas IE2 predominantly elicits CD4⁺ T cell responses, IE1 drives strong *ex vivo* CD8⁺ T cell reactivity^38^. These data would argue that an optimal SLP vaccine targeting the early phase of HCMV infection might require a combination of IE1 and IE2 peptides to achieve balanced CD4⁺ and CD8⁺ T cell immunity. Notably, SLP3 is shared between IE1 and IE2; whether differential processing enhances protective efficacy warrants further investigation.

The development of effective immunotherapies against HCMV remains a major research priority. To date, IE2 has not been incorporated into antibody- or T cell-based vaccine platforms. Our findings demonstrate that IE2 elicits highly polyfunctional CD4⁺ T cell responses detectable *ex vivo* and that its immunogenicity is comparable to pp65. The identification of highly immunogenic IE2 T cell epitopes provide a valuable resource for future vaccine design and adoptive T cell therapies targeting HCMV.

## Data availability

The dataset comprises large flow cytometry files and associated metadata deposited at Leiden University Medical Center (LUMC) and is available upon reasonable request from the corresponding authors.

## Acknowledgements

We also thank Ierry-Ann Lourens and Ilina Ehsan for their valuable contributions and technical assistance with the experiments. We thank Dr. Jan Wouter Drijfhout and the GMP-Peptide Facility at LUMC for peptide synthesis.

## Authors’ Contribution

EP, MJPW, RA, and SHvdB designed the research. EP performed the experiments and analyzed the data. EP wrote the first draft of the manuscript. RA and SHvdB contributed to manuscript writing and editing.

## Figure legends

**Supplementary Figure 1.**
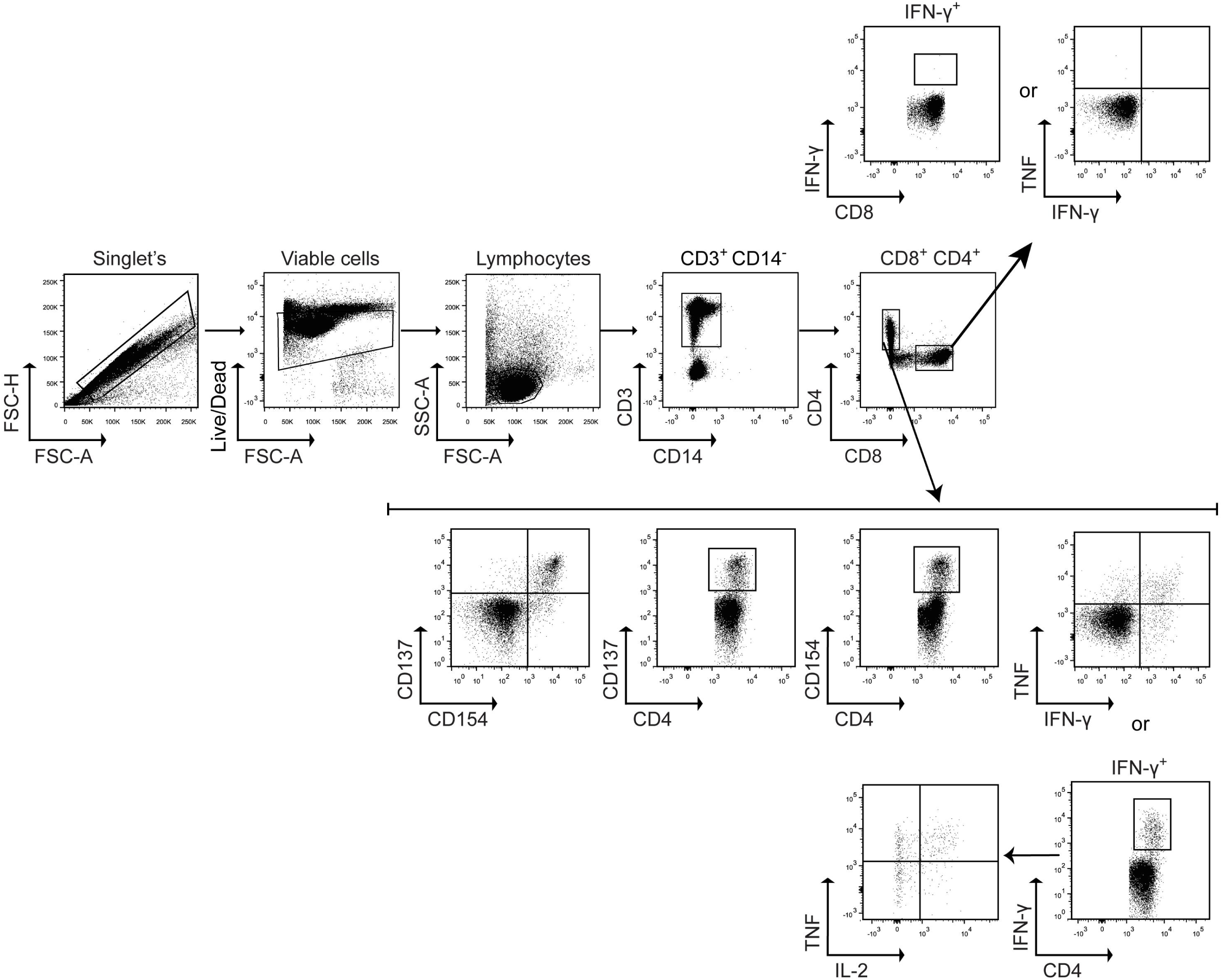
Gating strategy for intracellular cytokine staining of IE2-specific CD4⁺ and CD8⁺ T cells. Cells were sequentially gated for singlets (SSC-A vs. FSC-H), viability (FSC-A vs. live/dead), and lymphocytes (FSC-A vs. SSC-A). Lymphocytes were then gated on CD3⁺CD14⁻ to exclude monocytes and analyzed for IFN-γ⁺ CD4⁺ and CD8⁺ T cells. These subsets were further assessed for TNF and IL-2 production, as well as expression of the activation markers CD154 and CD137.

**Supplementary Figure 2:**
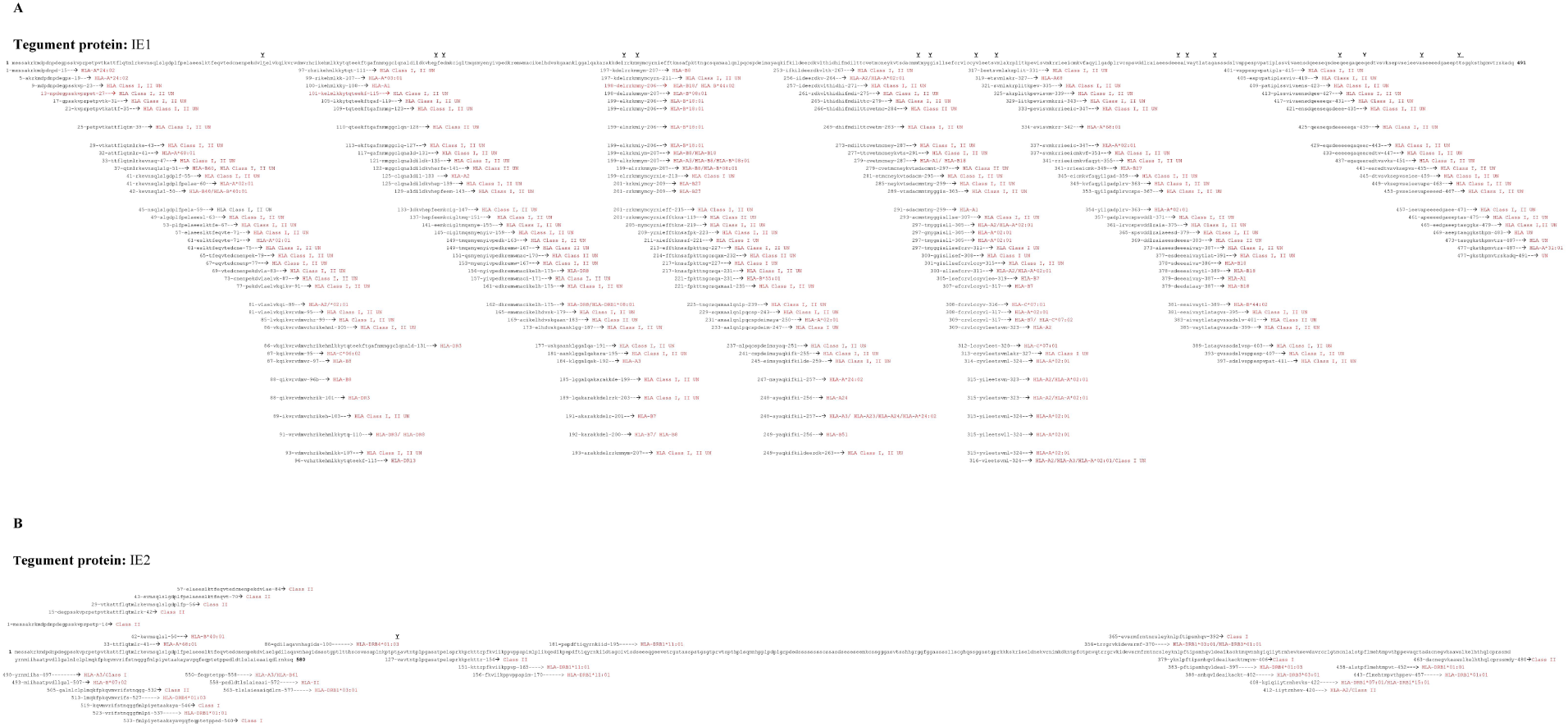
Chart of all known IE1 and IE2 HCMV T cell epitopes.

## Tables

**Supplementary Table 1.** IE2 peptide pools and individual SLPs used in this study. All synthetic long peptides (SLPs) covering the HCMV IE2 protein are listed, including amino acid positions and sequences. Eight peptide pools, each comprising 5-6 overlapping SLPs, are indicated. Individual SLPs contained within each pool are shown, providing a complete overview of the peptides used for *ex vivo* and *in vitro* T cell assays.

|  | SLP # | Peptide residues (aa) | SLP Sequence |
| --- | --- | --- | --- |
| pool 1 | 1 | IE2<sub>1-14</sub> | MESSAKRKMDPDNPDEGPSSKVP RPETP |
|  | 2 | IE2<sub>15-42</sub> | DEGPSSKVP RPETPVTKATTF LQTMLRK |
|  | 3 | IE2<sub>29-56</sub> | VT KATTF LQTMLRKEVNSQLSLGDPLFP |
|  | 4 | IE2<sub>43-70</sub> | EVNSQLSLGDPLFP ELAEESLKT FEQVT |
|  | 5 | IE2<sub>57-84</sub> | ELAEESLKT FEQVTEDCNENPEKDV LAE |
| pool 2 | 6 | IE2<sub>71-98</sub> | EDCNENPEKDV LAELGDILAQAVNHAGI |
|  | 7 | IE2<sub>85-112</sub> | LG DILAQAVNHAGIDSSSTGPTLTTHSC |
|  | 8 | IE2<sub>99-126</sub> | DSSSTGPTLTTHSCSVSSAPLNKPTPTS |
|  | 9 | IE2<sub>113-140</sub> | SVSSAPLNKPTPTSVAVTNTPLPGASAT |
|  | 10 | IE2<sub>127-154</sub> | VAVTNTPLPGASATPELSPRKKPRKTTR |
| pool 3 | 11 | IE2<sub>141-168</sub> | PELSPRKKPRKTTRPFKVIKPPVPPAP |
|  | 12 | IE2<sub>154-182</sub> | PFKVIKPPVPPAPIMLPLIKQEDIKPE |
|  | 13 | IE2<sub>169-196</sub> | IMLPLIKQEDIKPEPDFTIQYRNKIIDT |
|  | 14 | IE2<sub>183-210</sub> | PDFTIQYRNKIIDTAGCIVISDSEEEQG |
|  | 15 | IE2<sub>197-224</sub> | AGCIVISDSEEEQGEVETR GATASSPS |
| pool 4 | 16 | IE2<sub>211-238</sub> | EEVETR GATASSPSTGSGT PRVTSPTHP |
|  | 17 | IE2<sub>225-252</sub> | TGSGT PRVTSPTHP LSQLMNHPLPDPLG |
|  | 18 | IE2<sub>239-266</sub> | LSQMNHPPLPDPLGRPDE DSSSSSSSSC |
|  | 19 | IE2<sub>253-280</sub> | RPDE DSSSSSSSSC SSASDSESESEEMK |
|  | 20 | IE2<sub>267-294</sub> | SSASDSESESEEMKCSSGGGASVTSSH |
| pool 5 | 21 | IE2<sub>281-308</sub> | CSSGGGASVTSSHGRGGFGGAASSLL |
|  | 22 | IE2<sub>295-322</sub> | GRGGFGGAASSLLSCGHQSSGGASTGP |
|  | 23 | IE2<sub>309-336</sub> | SCGHQSSGGASTGPRKKSKRIELDNE |
|  | 24 | IE2<sub>323-350</sub> | RKKKSKRIELDNEKVRNIMKDKNTPFC |
|  | 25 | IE2<sub>337-364</sub> | KVRNIMKDKNTPFCTPNVQTRGRVKID |
| pool 6 | 26 | IE2<sub>351-378</sub> | TPNVQTRGRVKIDEVSRMFRNTNRSLE |
|  | 27 | IE2<sub>365-392</sub> | EVSRMFRNTNRSLEYKNLPFTIPSMHQV |
|  | 28 | IE2<sub>379-406</sub> | YKNLPFTIPSMHQVLDEAIKACKTMQVN |
|  | 29 | IE2<sub>393-420</sub> | LDEAIKACKTMQVNNKGIQIYTRNHEV |
|  | 30 | IE2<sub>407-434</sub> | NKGIQIYTRNHEVKSEVDAVRCRLGTM |
| pool 7 | 31 | IE2<sub>421-448</sub> | KSEVDAVRCRLGTM CNLALSTPFLMEHT |
|  | 32 | IE2<sub>435-462</sub> | CNLALSTPFLMEHTMPVTHPPEVAQRTA |
|  | 33 | IE2<sub>449-476</sub> | MPVTHPPEVAQRTADACNEG VKAAWSLK |
|  | 34 | IE2<sub>463-480</sub> | DACNEG VKAAWSLKELHTHQLCPRSSDY |
|  | 35 | IE2<sub>477-504</sub> | ELHTHQLCPRSSDYRNMIHAATPVDLL |
| pool 8 | 36 | IE2<sub>481-518</sub> | RNMIHAATPVDLLGALNLCLPLMQKFP |
|  | 37 | IE2<sub>505-532</sub> | GALNLCLPLMQKFPKQVMVRIFSTNQGG |
|  | 38 | IE2<sub>519-546</sub> | KQVMVRIFSTNQGGFMLPIYETAAKAYA |
|  | 39 | IE2<sub>533-560</sub> | FMLPIYETAAKAYAVGQFEQPTETPPED |
|  | 40 | IE2<sub>547-574</sub> | VGQFEQPTETPPEDLDTLSLAIEAAIQD |
|  | 41 | IE2<sub>561-580</sub> | LDTLSLAIEAAIQDLRNKSQ |

**Supplementary Table 2.**
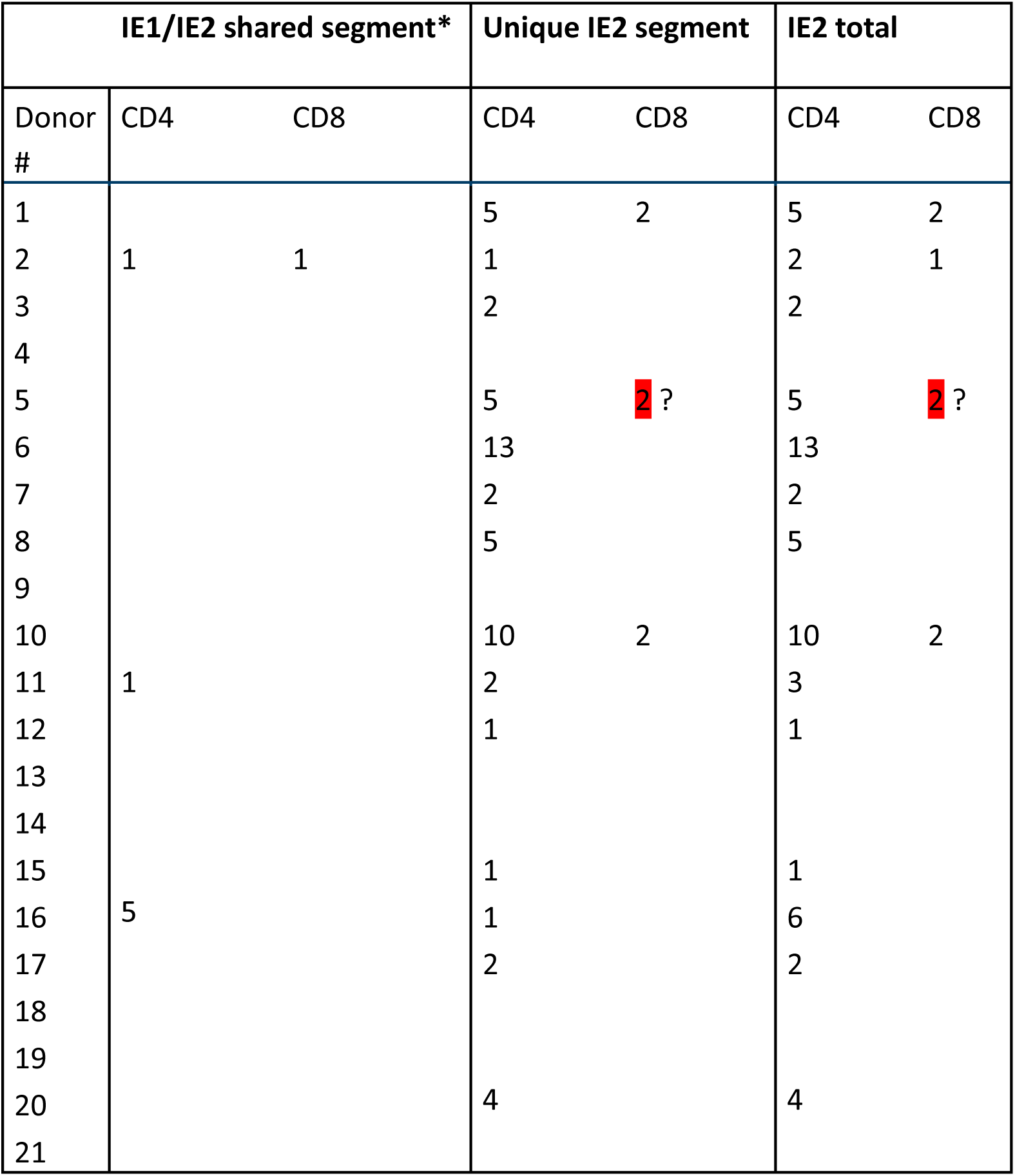
IE1- and IE2-specific CD4⁺ and CD8⁺ T cell responses per donor. Twenty-one donors were assessed for CD4⁺ and CD8⁺ T cell recognition of 28-mer IE2 peptides by ICS. Because the first 85 amino acids of IE1 and IE2 are identical, responses are divided into three segments: shared IE1/IE2 and unique IE2. The total number of single and shared T cell responses is shown. CD4⁺ responses were confirmed in 11 donors, including three CMV-seronegative donors (16, 17, 20), whereas CD8⁺ responses were detected in three CMV-seropositive donors.

| Donor # | IE1/IE2 shared segment* |  | Unique IE2 segment |  | IE2 total |  |
| --- | --- | --- | --- | --- | --- | --- |
|  | CD4 | CD8 | CD4 | CD8 | CD4 | CD8 |
| 1 |  |  | 5 | 2 | 5 | 2 |
| 2 | 1 | 1 | 1 |  | 2 | 1 |
| 3 |  |  | 2 |  | 2 |  |
| 4 |  |  |  |  |  |  |
| 5 |  |  | 5 | 2 ? | 5 | 2 ? |
| 6 |  |  | 13 |  | 13 |  |
| 7 |  |  | 2 |  | 2 |  |
| 8 |  |  | 5 |  | 5 |  |
| 9 |  |  |  |  |  |  |
| 10 |  |  | 10 | 2 | 10 | 2 |
| 11 | 1 |  | 2 |  | 3 |  |
| 12 |  |  | 1 |  | 1 |  |
| 13 |  |  |  |  |  |  |
| 14 |  |  |  |  |  |  |
| 15 |  |  | 1 |  | 1 |  |
| 16 | 5 |  | 1 |  | 6 |  |
| 17 |  |  | 2 |  | 2 |  |
| 18 |  |  |  |  |  |  |
| 19 |  |  |  |  |  |  |
| 20 |  |  | 4 |  | 4 |  |
| 21 |  |  |  |  |  |  |

---

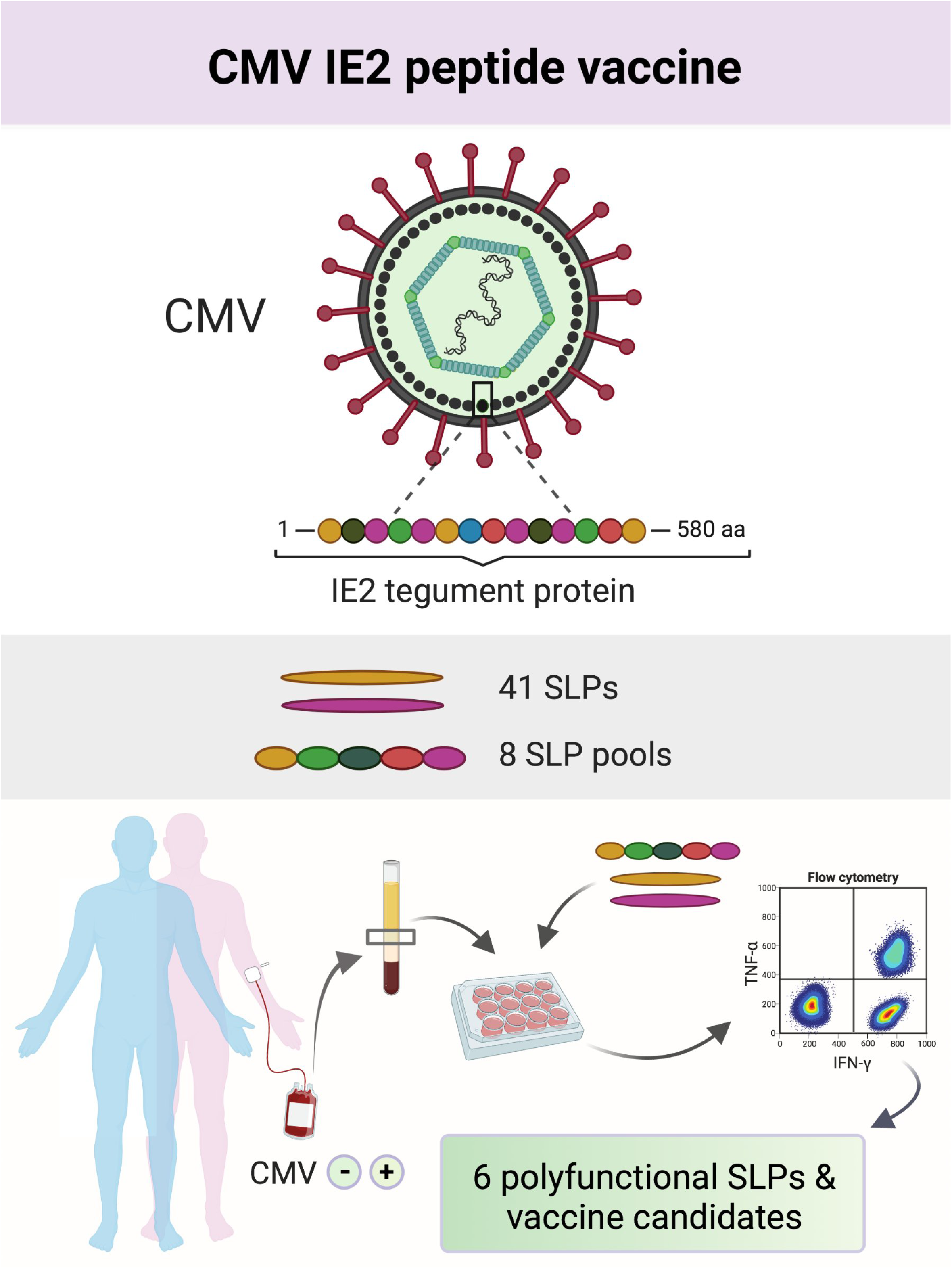

